# PeptiVerse: A Unified Platform for Therapeutic Peptide Property Prediction

**DOI:** 10.64898/2025.12.31.697180

**Authors:** Yinuo Zhang, Sophia Tang, Tong Chen, Elizabeth Mahood, Sophia Vincoff, Pranam Chatterjee

## Abstract

Therapeutic peptides combine the advantages of small molecules and antibodies, offering target flexibility and low immunogenicity, yet their successful translation requires careful evaluation of multiple developability properties beyond binding alone. As chemically modified peptides become increasingly common in drug design, no unified platform currently supports systematic property assessment across both canonical sequences and SMILES-based representations. Leveraging the generalizability of large foundational models trained on protein and chemical data, we introduce **PeptiVerse**, a universal therapeutic peptide property prediction platform. PeptiVerse accepts either amino acid sequences or chemically modified peptide SMILES, delivers state-of-the-art performance across diverse property prediction tasks, and provides both a web interface and open-source implementation for rapid, accessible, and scalable peptide developability analysis. By unifying property prediction across representations, PeptiVerse directly supports early-stage peptide therapeutic development campaigns and property-aware generative design workflows.

## 1 Introduction

Peptide-based therapeutics have gained significant attention in recent years, highlighted by the clinical success of GLP-1 receptor agonists for metabolic diseases [Zheng et al., 2024, Drucker, 2025]. As a therapeutic modality, peptides occupy a unique position between small molecules and antibodies, combining larger interaction surfaces capable of engaging protein-protein interfaces traditionally considered undruggable [Chen et al., 2025a, Wang et al., 2022] with reduced immunogenicity and manufacturing complexity relative to full-length antibodies [Wang et al., 2022, Tang et al., 2025a, Chen et al., 2023]. These features make peptides candidates for a broad spectrum of therapeutic targets, including receptors, enzymes, and intrinsically disordered regions.

Despite these advantages, native peptides often exhibit suboptimal translational profiles. For example, poor membrane permeability limits cellular uptake and oral bioavailability [Storchmannová et al., 2025, Chen et al., 2022], rapid proteolytic degradation results in short circulating half-life and frequent dosing [Werle and Bernkop-Schnürch, 2006, Wang et al., 2022], and low solubility promotes aggregation and formulation failure [Malavolta et al., 2006, Zapadka et al., 2017]. In addition, certain features of amphipathic sequences can cause hemolysis through nonspecific membrane disruption [Kellermeyer, 1962, Timmons and Hewage, 2020], while nonspecific protein adsorption (fouling) reduces effective concentration and increases off-target interactions in complex biological environments [Jiang and Cao, 2010, Guntuboina et al., 2023]. These limitations can be partially mitigated through cyclization, terminal modifications, D-amino acids, or other noncanonical residues [Wang et al., 2022, Li et al., 2023, DeGruyter et al., 2017], but such interventions push peptides beyond the assumptions of traditional protein sequence-based predictors [Notin et al., 2023, Xie et al., 2025]. As a result, practical peptide design requires systematic evaluation of multiple experimentally grounded properties beyond binding affinity alone.

Existing computational tools inadequately address this need. Sequence-based predictors such as Peptide-BERT are confined to natural amino acids and cannot accommodate chemical modifications [Guntuboina et al., 2023]. Small-molecule ADMET platforms accept “chemical” inputs in the form of SMILES but are trained on drug-like chemical space that differs substantially from peptides and proteins [Fu et al., 2024, Swanson et al., 2024]. Peptide-oriented SMILES predictors, including PepLand, PepDoRA, and PeptideDashboard, represent important steps toward chemistry-aware peptide modeling [Zhang et al., 2025a, Wang et al., 2024, Ansari and White, 2023], yet cover only a limited subset of relevant properties. Fitness-focused platforms such as ProteinGym benchmark mutational effects but do not target critical biochemical traits relevant for therapeutic peptide design [Notin et al., 2023, Xie et al., 2025]. Together, these gaps highlight the lack of a comprehensive, modality-flexible predictor capable of evaluating the full chemical and sequence diversity of modern peptide therapeutics.

The recent emergence of large protein and chemical language models enables unified property predictors that leverage rich learned representations without requiring explicit structural information [Lin et al., 2023, Feller and Wilke, 2025a]. Such predictors have become integral to modern generative peptide design workflows, where classifiers guide sampling, rank candidates, or provide post hoc filtering [Chen et al., 2025b,c,d, Goel et al., 2025, Tang et al., 2025b,c,d, Zhang et al., 2025b, Vincoff et al., 2025]. In this setting, gradient coupling between generator and predictor is not always necessary, allowing effective use of classical methods such as SVM [Hearst et al., 1998], Elastic Nets [Simoncini, 2005], and XGBoost [Chen et al., 2015], which perform well on pretrained embeddings while reducing overfitting risk [De Landsheere et al., 2025].

Here, we introduce **PeptiVerse** (Figure 1), a universal therapeutic peptide property evaluation framework designed to standardize and accelerate computational peptide design. PeptiVerse integrates state-of-the-art foundational models with carefully curated datasets to deliver fast, accurate, and scalable property predictions, supporting both sequence-based and SMILES-based peptide inputs. Beyond post hoc evaluation, these predictors can be directly used as guidance oracles within generative modeling workflows [Tang et al., 2025b,c,d, Chen et al., 2025c,d], enabling generation, ranking, and optimization of peptide candidates across diverse targets. Together, PeptiVerse provides a unified foundation for property-aware peptide discovery, enabling both early-stage candidate prioritization and integration with generative design workflows to accelerate therapeutic translation.

**Figure 1:**
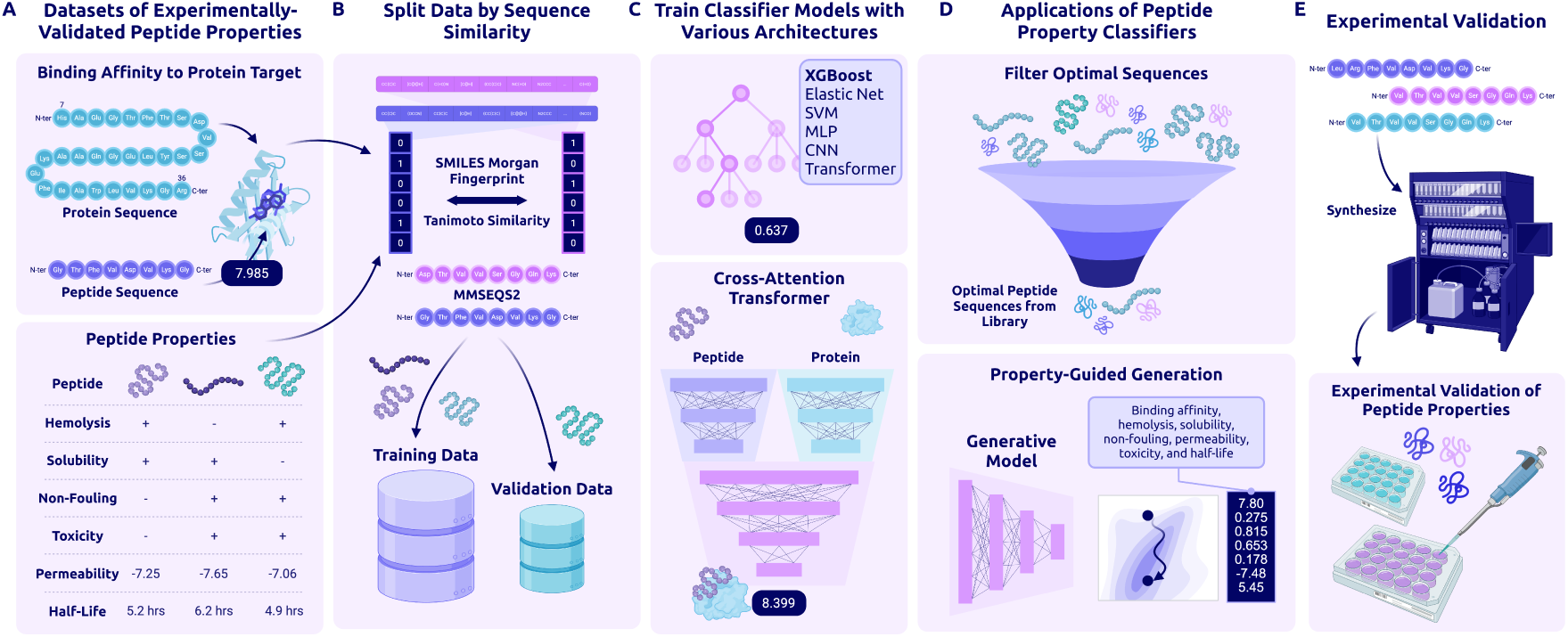
Overview of PeptiVerse Workflow and Applications. **(A)** Dataset collection: We collected datasets of experimentally-derived measurements for the following peptide properties: hemolysis, solubility, non-fouling, toxicity, permeability, half-life, and binding affinity to protein target(s). Measurements for all properties were collected across both canonical and non-canonical peptides. **(B)** Data splitting: Non-canonical peptides, represented by SMILES sequences, were split via Tanimoto similarity of Morgan fingerprints. Canonical peptide sequences were split via MMseqs2. **(C)** Property prediction: For most properties, we compared performance across multiple architectures, including XGBoost, Elastic Net, support vector machine (SVM), multi-layer perceptron (MLP), convolutional neural network (CNN), and Transformer. The binding affinity regression model is a cross-attention transformer operating upon the peptide and protein sequence representations. **(D)** Applications: PeptiVerse predictions can be used to filter existing peptides for desired properties or as a reward signal for property-guided generative models. **(E)** Validation: Each predicted property can be validated with downstream wet-lab experiments.

## 2 Results

### 2.1 Dataset composition highlights constraints across peptide properties

Accurate peptide property prediction is fundamentally constrained by data availability, coverage, and experimental diversity. As an initial step, we curated and integrated experimentally derived datasets spanning multiple peptide properties from a wide range of public sources [Guntuboina et al., 2023, Zhang et al., 2025a, Li et al., 2023, Pirtskhalava et al., 2021, Rathore et al., 2024, Smialowski et al., 2012, Jain et al., 2024, Mathur et al., 2016, D’Aloisio et al., 2021]. These datasets collectively cover both canonical amino acid sequences and chemically modified peptides represented as SMILES, enabling unified analysis across representation modalities. Examination of dataset composition reveals substantial heterogeneity in dataset size, label balance, and value distributions across properties (Figure 2; Table 2).

**Figure 2:**
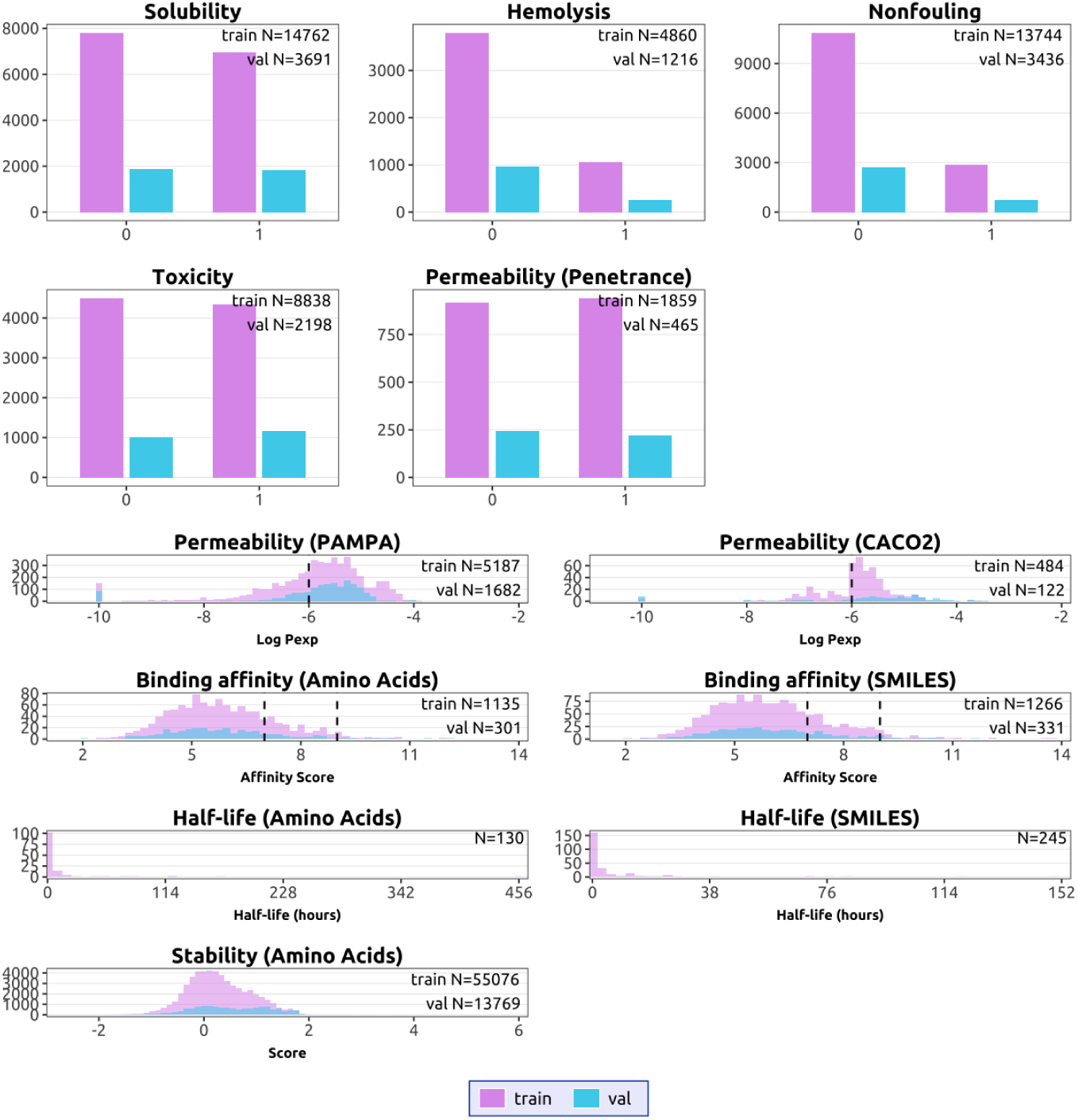
Training data distributions. Top: Distributions of binary-labeled datasets across training and validation splits. Bottom: Distributions of continuous-valued datasets across training and validation splits. Purple denotes training samples and blue denotes validation samples. Dashed vertical lines indicate heuristic thresholds for favorable properties, including good permeability (*x* = *−*6) and strong binding affinity (*x* = 7 and 9). Due to limited data availability, half-life predictors were trained using cross-validation and therefore are shown without an explicit train/-validation split.

**Table 1:** Results for property prediction benchmarks. The results reported for PepLand are taken directly from the original publication [Zhang et al., 2025a], which uses random data splits, whereas PeptiVerse employs similarity-aware splits to evaluate out-of-distribution generalization. AA denotes amino acid sequence inputs.

| Method | Regression (Spearman Correlation) |  |  | Classification (Best F1) |  |
| --- | --- | --- | --- | --- | --- |
|  | Binding Affinity (AA) | Permeability (all SMILES combined) | Binding Affinity (SMILES) | Permeability (Penetrance) | Solubility |
| ESM-2 | 0.452 | – | – | 0.885 | 0.725 |
| PepLand | 0.503 | 0.628 | **0.768** | 0.838 | 0.662 |
| PepLand + ESM-2 | 0.454 | – | – | 0.885 | 0.730 |
| PeptiVerse | **0.557** | **0.671** | 0.582 | **0.929** | **0.754** |

**Table 2:**
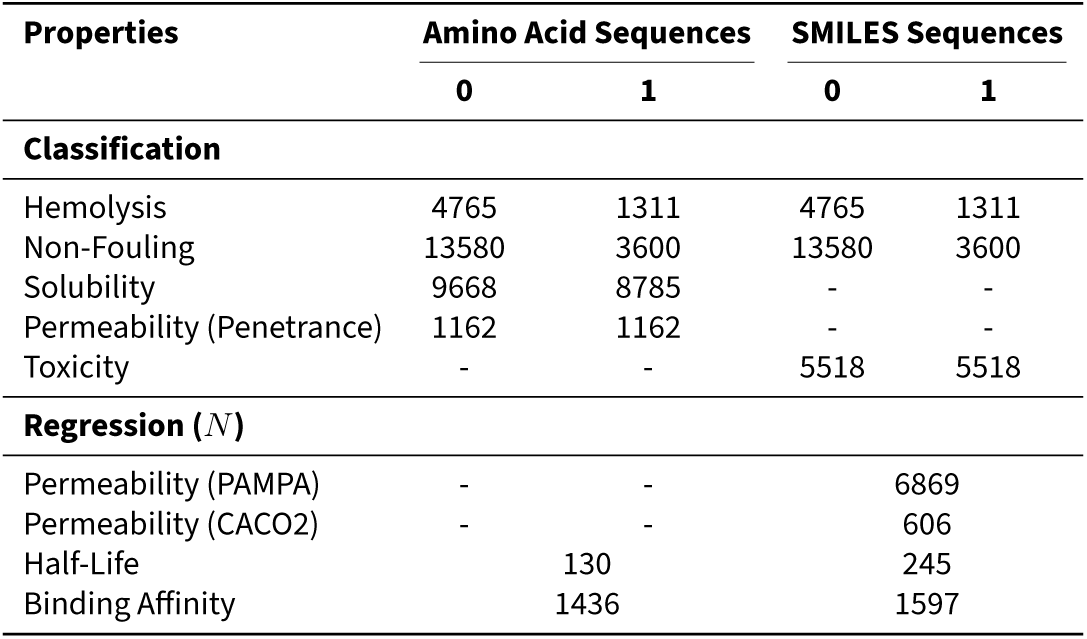
Dataset composition across peptide property prediction tasks. Classification tasks report the number of samples per class (0/1), where 0 is negative, and 1 is positive. Regression tasks report the total number of samples (*N* ). Counts are shown separately for amino acid and SMILES sequence representations, illustrating differences in data availability and class balance across properties.

Our data show that several classification tasks are supported by large, well-populated datasets with broad coverage of peptide sequence and chemical space. Hemolysis and non-fouling datasets comprise thousands to tens of thousands of peptides curated from antimicrobial and surface-interaction studies [Pirtskhalava et al., 2021, Barrett et al., 2018], while solubility datasets aggregate protein expression outcomes from structural genomics pipelines and mutational databases [Smialowski et al., 2012, Velecký et al., 2022]. Permeability datasets include both canonical cell-penetrating peptides and noncanonical cyclic peptides measured using PAMPA and Caco-2 assays [Avdeef, 2005, Artursson et al., 2012, Li et al., 2023], yielding relatively balanced class distributions and continuous-valued measurements spanning multiple orders of magnitude. These properties exhibit broad value distributions and sufficient sample sizes to support robust model training and evaluation under similarity-aware splits.

In contrast, regression tasks such as peptide half-life and binding affinity remain comparatively data-limited. Half-life measurements, curated from THPdb2, PEPlife, and PepTherDia [Jain et al., 2024, Mathur et al., 2016, D’Aloisio et al., 2021], are sparse, heterogeneous in experimental protocol, and often reported in coarse or qualitative units, resulting in limited sample sizes for both sequence- and SMILES-based representations. Binding affinity datasets aggregate diverse experimental readouts (*K_d_*, *K_i_*, and *IC*_50_) across protein–peptide pairs [Zhang et al., 2025a, Lei et al., 2021], but remain modest in scale relative to classification tasks. Together, our observations highlight that achievable predictive performance across peptide properties is frequently constrained by data availability and experimental variability rather than model capacity alone, motivating property-specific modeling strategies and emphasizing the importance of continued dataset expansion.

### 2.2 PeptiVerse deploys state-of-the-art predictors for a broad range of therapeutically relevant peptide properties

Given our heterogeneous data settings, we evaluated a diverse set of predictor architectures within PeptiVerse, including linear models, boosting methods, multilayer perceptrons (MLPs), convolutional neural networks (CNNs), support vector machines (SVMs), and transformer-based models (Figure 3). Across all classification tasks and both input modalities for amino acid sequence inputs and SMILES inputs, overall performance differences between architectures were modest when trained on fixed, information-rich embeddings derived from ESM-2 and PeptideCLM (Supplementary Figure S1; Supplementary Table S1). This consistency suggests that representation quality and dataset characteristics, rather than downstream model capacity, constitute the primary performance bottleneck in peptide property prediction.

**Figure 3:**
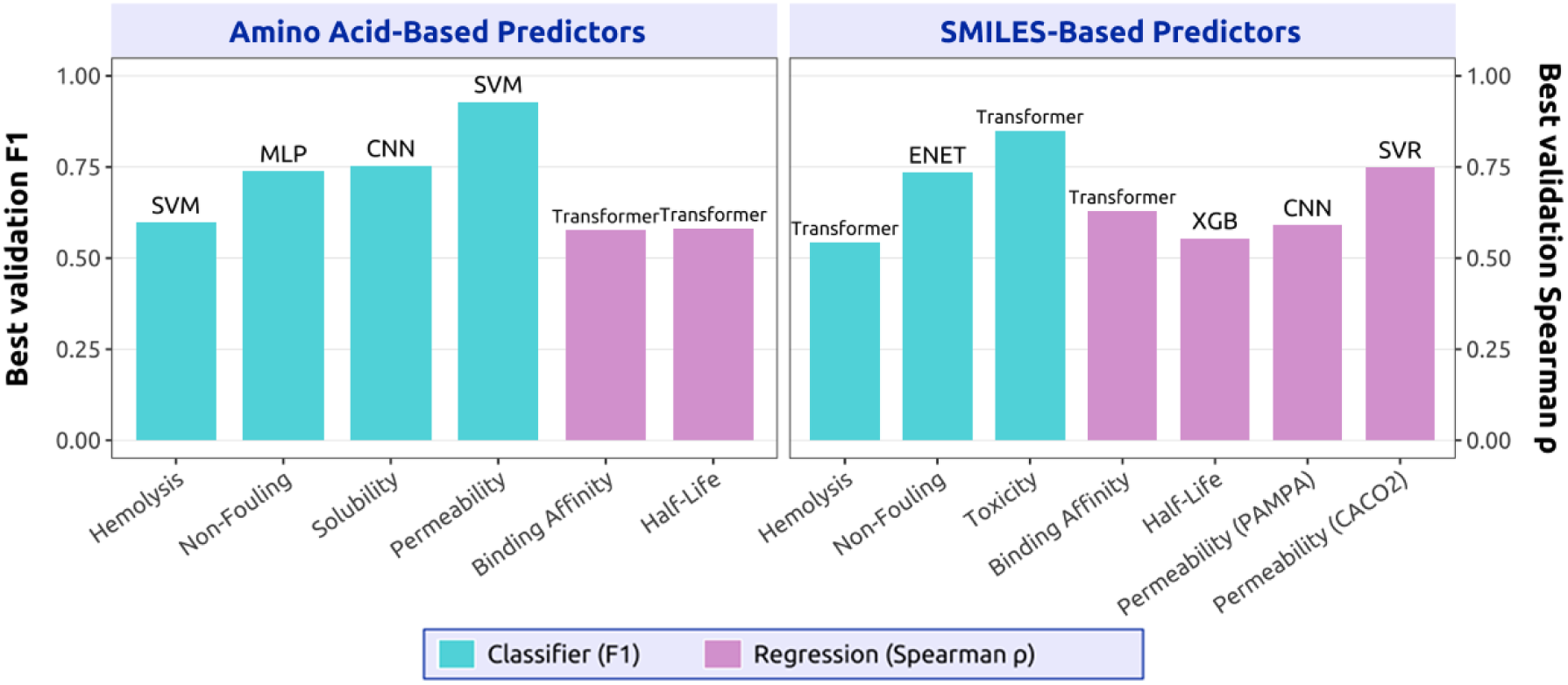
Best-model performance of PeptiVerse across different therapeutic peptide properties. Left: Performance on amino acid encoded sequences using ESM-2 embeddings; Right: Performance on SMILES represented sequences using PeptideCLM embeddings. Bars report validation set performance of the best-performing model selected after OPTUNA optimization, with names of the model annotated on top. Cyan indicates the performance of the classification models, and light purple indicates the performance of the regression models.

Rather than enforcing a single architecture, PeptiVerse exposes the best-performing model per property and representation, reflecting the observation that no single architecture emerges as universally optimal. Different peptide properties favored different inductive biases, with hemolysis and permeability benefiting from margin-based or kernel methods, solubility favoring convolutional architectures, and chemically rich SMILES-based tasks often selecting transformer-based heads (Figure 3). Importantly, similar trends were observed for regression tasks, where multiple nonlinear models achieved comparable performance, and no clear architectural dominance was observed (Supplementary Figure S2; Supplementary Figure S3). In both settings, properties supported by larger datasets with broader coverage of the underlying value distributions, such as permeability, consistently yielded stronger performance than properties with limited or sparsely distributed data, such as peptide half-life.

Despite this diversity, tree-based boosting models such as XGBoost consistently achieve competitive performance across tasks while offering advantages in training stability, computational efficiency, and ease of deployment. This balance makes lightweight statistical models particularly attractive for large-scale screening and integration into generative design pipelines, especially when combined with strong pretrained embeddings. Importantly, all evaluations were conducted using similarity-based data splits, indicating that simple models can generalize comparably to deep architectures on out-of-distribution peptide designs when initialized with high-quality representations.

### 2.3 PeptiVerse provides experimentally-meaningful binding affinity predictions in contrast to structure-based predictors

To assess whether PeptiVerse binding affinity predictions reflect physically meaningful interaction strength, we examined their relationship to structure-based confidence metrics derived from state-of-the-art *de novo* protein-peptide complex predictors. Recent studies have reported correlations between structure prediction confidence scores, such as ipTM, and docking-based interactions metrics for protein-protein interactions, motivating the use of such scores as proxies for binding strength in *ab initio* modeling pipelines [Zhu et al., 2023, Peng et al., 2025]. We therefore asked whether similar relationships hold for peptide-protein interactions, including both wild-type and chemically modified peptides.

Using OpenFold3-predicted complexes [The OpenFold3 Team, 2025], we compared ipTM scores against experimentally measured binding affinities for 1,436 amino acid sequence inputs and 1,597 SMILES-based peptide inputs. In contrast to prior observations for protein–protein interactions, ipTM showed negligible association with experimental binding affinity across either peptide representation (Supplementary Figure S4; |*ρ*| *≈* 0.05), indicating that structure confidence metrics are insufficient proxies for peptide–protein binding strength. While ipTM has proven effective for estimating protein–protein interaction strength, these results suggest that it does not reliably capture the energetic or kinetic determinants governing peptide binding, likely due to the increased flexibility, shallow binding interfaces, and diverse chemistries characteristic of peptide ligands. Moreover, even when accurate complex structures are available, structure prediction is computationally heavier than embedding-based inference and is not well-suited for high-throughput screening across large peptide libraries. PeptiVerse therefore provides a fast affinity surrogate that complements structure prediction rather than replacing it.

In contrast, PeptiVerse binding affinity predictions show statistically significant agreement with experimental measurements across both representation modalities, achieving Spearman *ρ* = 0.58 for amino acid sequence inputs and *ρ* = 0.56 for SMILES-based inputs (all *p <* 10*^−^*^3^; Supplementary Figure S3). PeptiVerse employs a transformer-based cross-attention architecture for binding affinity prediction that operates directly on protein and peptide embeddings. Unpooled embeddings provided a performance enhancement over pooled embeddings, supporting the hypothesis that preserving residue-level positional information is critical for modeling peptide–protein binding (Supplementary Figure S5). Together, these findings demonstrate that PeptiVerse provides experimentally relevant binding affinity estimates that complement structural modeling and enable binding-aware filtering and generative peptide design.

### 2.4 PeptiVerse demonstrates superior predictive performance relative to existing peptide property predictors

PeptiVerse is, to our knowledge, the first unified framework capable of predicting multiple physicochemical and developability properties for both amino acid sequence inputs and SMILES-encoded peptide inputs. In contrast, prior tools have made meaningful progress but remain specialized in either input modality or property scope [Feller and Wilke, 2025a, Zhang et al., 2025a]. PeptideBERT [Guntuboina et al., 2023] operates primarily on canonical amino acid sequences and does not generalize to chemically modified peptides. None of the existing tools provides the breadth of properties, multimodality, or user accessibility offered by PeptiVerse.

To quantitatively assess these differences, we compare PeptiVerse against representative prior methods across multiple regression and classification benchmarks in Table 1. Across most evaluated tasks, PeptiVerse achieves competitive or superior performance under a unified evaluation protocol. We note that PepLand reports higher performance on SMILES-based binding affinity prediction [Zhang et al., 2025a]. However, these results were obtained using random data splits, which permit substantial overlap in sequence or chemical similarity between the training and test sets. In contrast, PeptiVerse employs similarity-based splits, resulting in a substantially more challenging and realistic evaluation of out-of-distribution generalization. Performance differences on SMILES-based binding affinity, therefore, reflect differences in the evaluation protocol rather than representational limitations.

Additionally, the PepLand+ESM-2 setting is reported only for canonical peptide tasks, and the mechanism for switching between SMILES and sequence inputs is not explicitly specified. By contrast, PeptiVerse provides an explicit and unified multimodal design, enabling consistent evaluation across both canonical and non-canonical peptide spaces.

### 2.5 PeptiVerse is deployed as a unified web interface for peptide evaluation

Finally, we developed an interactive web interface to make PeptiVerse readily accessible for practical use by both experimental and computational researchers. As such, PeptiVerse is deployed as a web server (Figure 4) built with `Gradio` [Abid et al., 2019] and hosted on HuggingFace Spaces [Jain, 2022]. The interface allows users to submit peptide inputs as either amino acid sequences or SMILES strings, enabling property evaluation across both canonical and chemically modified peptides. A broad set of therapeutically relevant properties is supported, including binding affinity to a protein sequence target, hemolytic activity, non-fouling behavior, permeability, solubility, toxicity, and peptide half-life.

**Figure 4:**
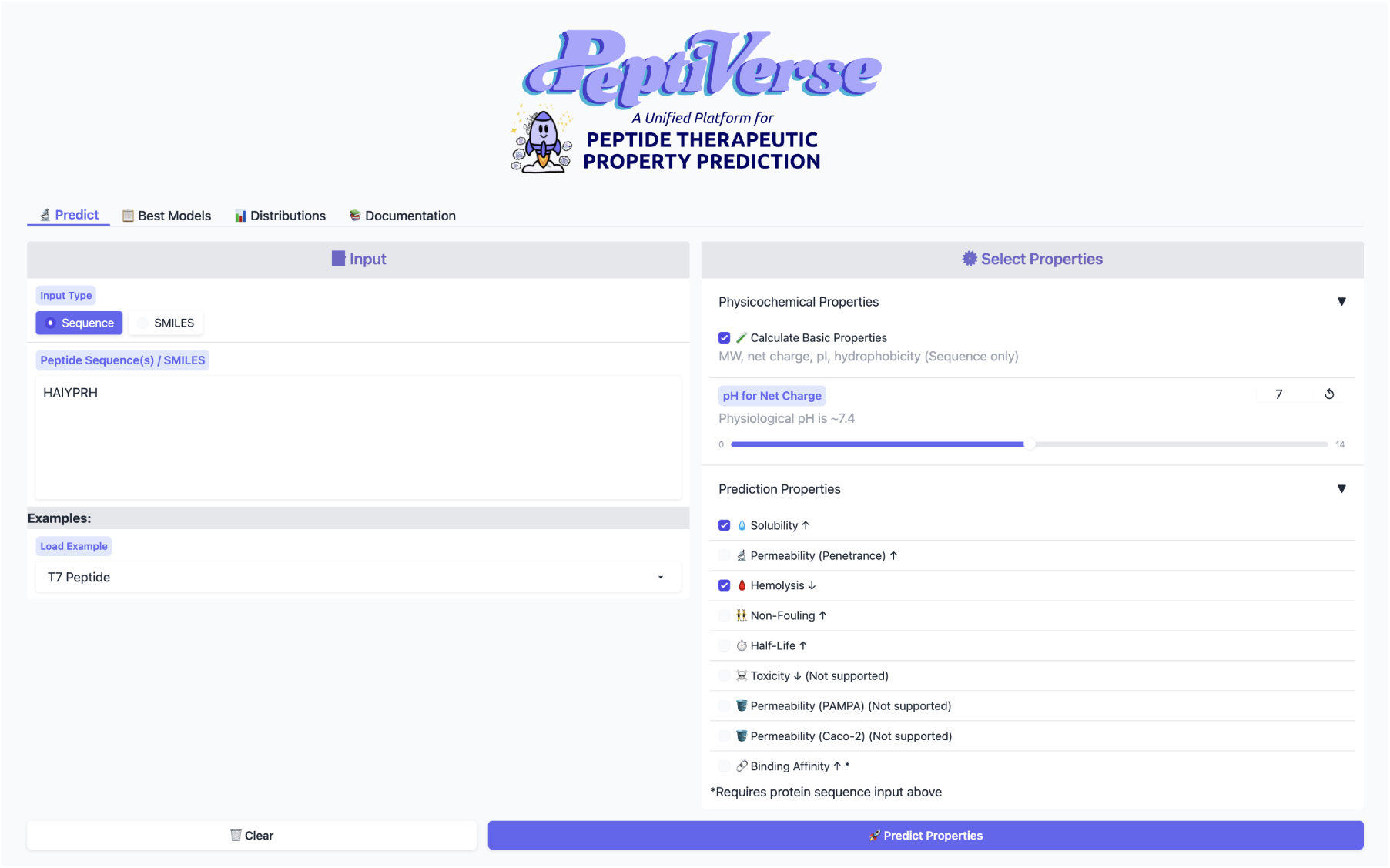
Web interface of PeptiVerse hosted on HuggingFace. The interface allows users to submit peptide inputs as either amino acid sequences or SMILES strings and select from a range of physicochemical and predictive properties for evaluation. Property options are dynamically enabled based on input type, and basic properties such as molecular weight, charge, and hydrophobicity can be computed alongside learned predictors. This design enables rapid, no-code peptide property assessment suitable for both exploratory analysis and integration into iterative design workflows.

To improve interpretability and usability, the interface provides visualizations of the underlying training data distributions, allowing users to contextualize their input peptides relative to experimentally characterized datasets. All datasets used to train the deployed models are standardized and distributed in HuggingFace Dataset format, facilitating reproducibility, benchmarking, and integration into downstream peptide design and optimization workflows.

## 3 Discussion

PeptiVerse introduces a unified framework for therapeutic peptide property prediction that supports both amino acid sequence inputs and SMILES representations, including peptides containing unnatural amino acids. By building on pretrained protein (ESM-2) and chemical (PeptideCLM) language models [Lin et al., 2023, Feller and Wilke, 2025a], PeptiVerse focuses on training lightweight predictor heads rather than full representation models, enabling efficient, scalable, and easily deployable property evaluation. These results highlight the heterogeneity of peptide property landscapes and motivate a flexible, property-aware modeling strategy rather than reliance on a single predictor.

Compared with prior SMILES-based peptide models such as PepLand [Zhang et al., 2025a], which rely on complex graph constructions and full-model retraining, PeptiVerse shows that foundational embeddings paired with simple, well-regularized classifiers are sufficient (and often superior) for practical peptide property prediction. This design reduces computational cost while improving generalizability, making PeptiVerse compatible with modern peptide design pipelines. The advantage is particularly clear for binding affinity, where structure prediction confidence alone fails to track peptide-protein binding strength, whereas PeptiVerse yields strong, statistically significant agreement with experimental measurements. Together, these results motivate fast, data-driven affinity predictors that complement rather than replace structural modeling, and position PeptiVerse as an open, extensible benchmark for therapeutic peptide discovery.

To ensure continued relevance, PeptiVerse will be updated regularly as new data and predictors become available. While several property models already achieve strong performance, others (i.e., chemically-modified peptide half-life in SMILES space) remain constrained by limited and heterogeneous experimental data. We therefore encourage open-source deposition of peptide property measurements from both academic and industrial efforts. In parallel, we are developing new specialized binding predictors, including peptide isoform-specificity [Vincoff et al., 2025], motif-specificity [Chen et al., 2025b], and metal-binding propensity [Zhang et al., 2025b], which will be incorporated as additional properties within the PeptiVerse framework. By providing a standardized, extensible, and openly accessible platform for integrating new data and models, PeptiVerse establishes a practical mechanism for expanding predictive scope, enabling property-guided generative design, and supporting the translation of next-generation peptide therapeutics.

## 4 Methods

### 4.1 Data Collection and Preparation

Throughout this work, we distinguish peptide inputs by representation modality rather than biological origin. “Amino acid” inputs refer to canonical sequence-based representations processed by protein language models, while “SMILES” inputs refer to chemistry-aware molecular representations, which may include both canonical and non-canonical peptides.

#### Hemolysis

Hemolysis data were retrieved from PeptideBert and peptide-dashboard [Guntuboina et al., 2023, Ansari and White, 2023], and cross-validated against the original experimental records in DBAASP v3.0 [Pirtskhalava et al., 2021]. Peptides labeled as 1 are considered hemolytic, whereas 0 denotes non-hemolytic activity. The final dataset comprised 4,765 hemolytic and 1,311 non-hemolytic peptide entries.

#### Permeability

All permeability annotations were obtained from PepLand [Zhang et al., 2025a], which sources experimental measurements from CycPeptMPDB [Li et al., 2023]. The *noncanonical* dataset contains 7,334 non-canonical peptides with reported permeability values measured using either PAMPA [Avdeef, 2005] or Caco-2 assays [Artursson et al., 2012]. Permeability is reported as log *P*_exp_, the logarithm of the effective permeability coefficient, which reflects the peptide’s lipophilicity and its ability to passively diffuse across lipid membranes.

PAMPA and Caco-2 assays quantify different biological processes, with PAMPA measuring passive membrane permeability and Caco-2 assays relating more closely to intestinal absorption potential [Avdeef, 2005, Ottaviani et al., 2006, Van Breemen and Li, 2005]. For this reason, the two assay types were handled separately during data preparation, not following PepLand’s combined training strategy. There were 6,869 PAMPA sequences and 606 Caco-2 sequences collected in the end. Following CycPeptMPDB conventions, peptides with log *P*_exp_ *≥ −*6.0 were labeled as high permeability, and the remaining peptides as weak permeability. [Li et al., 2023].

The canonical permeability dataset contains 1,162 cell-penetrating peptides and 1,162 non-penetrating peptides, curated from PepLand [Zhang et al., 2025a]. This dataset was constructed to balance peptide length distributions between positive and negative classes. Positive peptides were originally collected from 22 independent cell-penetrating peptide studies, while negative examples were sourced from UniProt. A notable fraction of the positive peptides (approximately 8.7%) exhibited low sequence complexity, defined as a ratio of peptide length to the number of unique amino acids greater than 5, compared to only 0.95% among non-penetrating peptides. Although these low-complexity sequences are likely engineered, they were retained, as they may encode informative features relevant to membrane permeability.

#### Non-Fouling

Non-fouling annotations were obtained from PeptideBERT and Peptide-Dashboard [Guntuboina et al., 2023, Ansari and White, 2023], both of which source data from the dataset of Barrett et al. [2018]. In this context, non-fouling peptides are defined as sequences that resist nonspecific protein adsorption, while fouling peptides permit such adsorption, which can lead to functional loss and reduced performance [Barrett et al., 2018, Banerjee et al., 2011]. Peptides labeled as 1 are classified as non-fouling, and those labeled as 0 are considered fouling. The curated dataset comprised 13,580 non-fouling and 3,600 fouling peptide entries.

#### Toxicity

Toxicity data were obtained from ToxinPred3.0, which provides canonical amino acid peptide sequences with experimentally validated toxicity labels [Rathore et al., 2024]. The dataset contains 5,518 toxic and 5,518 non-toxic peptides, where label 1 denotes toxic and 0 denotes non-toxic. Amino acid sequences were converted to SMILES representations using RDKit [Landrum et al., 2025]. Molecular redundancy was reduced by clustering peptides using Morgan fingerprints (radius 2, 2048 bits, including chirality) and RDKit’s Butina clustering algorithm with a similarity threshold of 0.6.

#### Solubility

Solubility labels follow the protocol established in PROSO II [Smialowski et al., 2012], as used in PeptideBERT [Guntuboina et al., 2023], with additional sequences incorporated from SoluProtMutDB [Velecký et al., 2022]. In PROSO II, the soluble class (1) was assigned based on experimental metadata recorded in pepcDB, following the Protein Structure Initiative (PSI) pipelines [Berman et al., 2009]. A sequence was labeled soluble once it reached the “Soluble” stage or any downstream stage. Additional soluble entries were derived from PDB records (up to 2010) annotated with expression from *E. coli*, as crystallographic and NMR structure determination requires a protein that has been purified in solution. The insoluble class (0) consisted of constructs that remained in the “not Soluble” state for at least eight months, based on the different pepcDB releases. This yields a combined dataset of 8,785 soluble and 9,668 negative sequences in total.

#### Binding Affinity

Binding affinity prediction was performed using paired protein sequences and peptide SMILES from the PepLand dataset [Zhang et al., 2025a], which aggregates different experimental measurements together (*K_d_, K_i_, IC*_50_). Their data collection protocol follows CAMP [Lei et al., 2021], which subsets data from RCSB PDB [Kouranov et al., 2006] and DrugBank entries containing the label of “peptide” [Knox et al., 2024]. All scores were transformed based on the negative logarithm of the original affinity data into a unified scale. A shared unit scale was roughly designed to indicate the strength of binding: with 9 indicating strong *nM* to *pM* binders; 7-9 indicating *nM* to *µM* medium binders, and *<* 7 indicating weak *µM* binders. The data split was based on affinity score distribution matching, ensuring both splits containing the similar distribution style of data. The final dataset contained 1,436 peptide–protein pairs with canonical peptide sequences (amino acid inputs) and 1,597 pairs with SMILES-based peptides containing noncanonical chemistry.

#### Stability

Stability data for amino acid sequence inputs were collected and organized from TAPE [Rao et al., 2019] and SaProt [Su et al., 2023] (Huggingface: `SaProtHub/Dataset-Stability-TAPE`) were used to pretrain stability predictors before half-life modeling. Stability was treated as a continuous score reflecting the ability of proteins to remain folded above concentration thresholds. A total of 68,845 sequences were collected.

#### Half-Life

Half-life data were compiled from THPdb2 [Jain et al., 2024], PEPlife [Mathur et al., 2016], and PepTherDia [D’Aloisio et al., 2021]. Only human serum measurements were retained. Reported values across datasets vary in units and granularity. All half-life measurements were therefore converted into hours for interpretability. Details of the data construction are provided in the Supplementary Information. The curated dataset comprised 130 amino acid sequence entries and 245 SMILES-based sequences.

#### Protein-peptide ipTM scores

The interface predicted TM-score (ipTM) is a confidence metric originally designed to assess the accuracy of predicted protein–protein interaction interfaces in multimeric structure prediction models [Abramson et al., 2024]. All ipTM scores for protein-peptide pairs, including SMILES represented pairs, were retrieved from API calling for the OpenFold3 [The OpenFold3 Team, 2025] through NVIDIA NGC platform [NVIDIA Corporation, 2025]. MSA profiles for the target proteins were constructed via MMseqs2 [Steinegger and Söding, 2017] against the Uniref30 database [Consortium, 2019].

#### Additional Properties

Additional physicochemical features, including isoelectric point, molecular weight, and hydrophobicity, were computed using utilities from `Biopython` package [Cock et al., 2009]. These properties were calculated dynamically with a tunable pH parameter to account for protonation-state dependence.

#### Data Splitting

All properties defined on amino acid–based peptide representations were split using a shared clustering strategy. Sequences from all amino acid sequence datasets were clustered independently using MMseqs2 [Steinegger and Söding, 2017] with identical parameters (`-min-seq-id 0.3 -c 0.8 -cov-mode 0`) and split using an 80/20 cluster-level split to prevent sequence leakage between training and evaluation subsets. Amino acid sequences were converted into SMILES format with `fasta2smi` command from `p2smi` [Feller and Wilke, 2025b]. For properties evaluated using both sequence and SMILES inputs (e.g., hemolysis, non-fouling), the original cluster-based train/validation assignments were preserved after conversion, ensuring fair comparison across input modalities. For datasets natively represented in SMILES space, molecules were clustered using Morgan fingerprints and Tanimoto distance via RDKit [Landrum et al., 2025], followed by an 80/20 cluster-level split. Binding affinity prediction, which involves paired peptide–protein inputs across two molecular modalities, was instead split by matching affinity score distributions between training and evaluation sets. Detailed dataset sizes, label distributions, and train/validation splits for all properties are summarized in Table 2 and Figure 2.

### 4.2 Model Architecture and Training

#### Sequence and SMILES Representations

Protein sequences were represented using embeddings from ESM-2 (`esm2_t33_650M_UR50D`) [Lin et al., 2023] while peptide SMILES were embedded via PeptideCLM (`PeptideCLM-23M-all`) [Feller and Wilke, 2025a], which employs a tokenizer of size 586 designed to capture noncanonical peptide chemistry. Depending on the downstream model, embeddings were either average-pooled across sequence positions to form fixed-length representations or retained in unpooled, position-resolved form to preserve residue-level and positional information. This distinction allowed different model classes to exploit either global sequence summaries or fine-grained positional structure.

#### Predictor Architectures

For properties including toxicity, solubility, hemolysis, non-fouling, and permeability, lightweight predictor heads were trained on frozen foundational embeddings. We evaluated a broad set of predictor architectures, including multilayer perceptrons (MLPs), convolutional neural networks (CNNs), and transformer-based models, as well as classical statistical learners such as Elastic Nets (ENET) [Zou and Hastie, 2005], support vector machines (SVMs) [Hearst et al., 1998], epsilon-support vector regression (SVR) [Carrasco et al., 2019], and XGBoost (XGB) [Chen et al., 2015]. ENET and SVM classifiers were implemented using the `RAPIDS cuML` library [Raschka et al., 2020] to enable GPU acceleration, while SVR models were implemented using `scikit-learn` [Pedregosa et al., 2011]. Classification models were trained using standard objective functions appropriate to each method, whereas regression models optimized mean squared error. All models were trained using their canonical formulations without architectural modification. For CNN- and transformer-based predictors, unpooled embeddings were used to allow convolutional kernels and self-attention mechanisms to explicitly exploit positional structure, while pooled embeddings were used for MLP and tree-based models operating on fixed-dimensional inputs.

#### Hyperparameter Optimization

Hyperparameters for all models were optimized using the Optuna framework [Akiba et al., 2019], with an initial target of 200 optimization trials per configuration (Supplementary Tables S2–S5). For parameters sampled on a logarithmic scale, values were drawn uniformly in log space to efficiently explore multiple orders of magnitude. For regression tasks, both mean squared error (MSE) and Huber loss were evaluated, using the latter to improve robustness to outliers via its *δ* parameter controlling the transition between L1 and L2 penalties. For computationally intensive configurations exceeding approximately 10 hours of wall-clock training time, the number of Optuna trials was reduced to 50 or 20 while preserving representative coverage of the hyperparameter space. Results of the hyperparameter exploration are reported in Supplementary Figures S2 and S6.

#### Binding Affinity Prediction

Binding affinity prediction was formulated using a transformer-based architecture with cross multi-head attention to learn a joint latent representation between peptide and protein modalities. The model architecture was fixed across experiments, with variation introduced only through peptide embedding initialization. Peptide inputs were represented using either ESM-2 or PeptideCLM embeddings [Lin et al., 2023, Feller and Wilke, 2025a], in pooled or unpooled form, while protein targets were consistently represented using ESM-2 embeddings. The model produced a continuous affinity score, which was additionally mapped to three discrete affinity classes, yielding a multitask formulation aligned with other property predictors. Hyperparameters were optimized using 200 Optuna trials, and Spearman’s rank correlation coefficient (*ρ*) was used as the primary model selection criterion (Supplementary Table S6; Supplementary Figure S2).

#### Half-Life Prediction

Peptide half-life prediction was formulated as a regression task using both amino acid sequence inputs and SMILES-based representations. For amino acid–based inputs, models were initialized from pre-trained stability predictors trained on the wild-type stability dataset and subsequently fine-tuned on half-life data. The same model families used for binary property prediction were retained. Four regression configurations were evaluated: (i) XGBoost predicting half-life in hours, (ii) XGBoost predicting log(1 + half-life), (iii) a transformer-based regressor predicting half-life in hours, and (iv) a transformer-based regressor predicting log(1 + half-life). Hyperparameters were optimized using Optuna with five-fold cross-validation, using 200 trials for XGB, SVR, and ENET models and 50 trials for CNN, MLP, and Transformer models, with 20 training epochs per trial.

For SMILES-based half-life prediction, models were trained directly on peptide SMILES embeddings without intermediate stability pretraining. Predictor architectures matched those used for amino acid–based half-life prediction and were trained to predict log(1 + half-life). Evaluation was performed using five-fold cross-validation. Optuna optimization employed 200 trials for XGB, SVR, and ENET models and 50 trials for CNN, MLP, and Transformer models, with transformer-based models trained for 100 epochs per trial.

#### Evaluation Metrics

Model evaluation metrics were selected according to task type. Classification performance was assessed using F1 score, area under the receiver operating characteristic curve (AUC), Matthews correlation coefficient (MCC), and accuracy, with results reported in Supplementary Figure S1 and Supplementary Table S1. Regression performance was evaluated using Pearson correlation coefficient (*r*), Spearman’s rank correlation coefficient (*ρ*), and the coefficient of determination (*R*^2^), with results summarized in Supplementary Figure S3.

## Declarations

### Data and Code Availability

The full set of predictors and datasets is available through a simple API at https://huggingface.co/ChatterjeeLab/PeptiVerse. Code for plotting in R is available at https://github.com/ynuozhang/Peptiverse_R.git. For users who prefer a no-code interface, all predictors can also be easily run via an interactive HuggingFace Space at https://huggingface.co/spaces/ChatterjeeLab/PeptiVerse. All code is open source, with predictors and associated resources updated and maintained regularly.

## Acknowledgments

We thank the entire experimental team of the Chatterjee Lab for productive input during the curation of these models and datasets. We also thank Lauren Hong for designing the PeptiVerse logo.

## Author Contributions

Y.Z., S.T., T.C., S.V., and E.M., trained models, curated datasets, and prepared results. Y.Z. managed dataset and model curation and wrote the paper, with input from S.T. and T.C. P.C. conceived, designed, and directed the study, and reviewed and finalized the manuscript.

## Supplementary Information

### Supplementary Methods

#### Half-Life Data Preparation

Peptide half-life can be strongly influenced by terminal modifications, D-amino acids, unnatural residues, and backbone cyclization [Mathur et al., 2016]. To preserve chemically relevant information, peptides were represented using SMILES whenever available, while sequence-based representations were also retained. For datasets lacking canonical SMILES, structures were generated from sequence-level information using the `fasta2smi` utility from `p2smi` [Feller and Wilke, 2025b]. Tokenized sequences with lengths over 200 were filtered out to avoid outliers.

Half-life entries from PepTherDia [D’Aloisio et al., 2021] were provided as canonical SMILES with qualitative or ranged lifespan descriptions. PepTherDia contains mostly either entries with unnatural amino acids or chemical modification at the ends, or sequences with cyclic connections. Entries such as “cannot be calculated” were excluded. Ranges such as “1–6 min” were approximated using the midpoint (taken as 900 seconds), and qualitative descriptions like “10 minutes or less” were converted to fixed values (600 seconds). THPdb2 [Jain et al., 2024] also reported half-life values using descriptive or numerical formats, but does not provide canonical SMILES. In addition, peptide sequences may omit D-amino acid annotations or represent noncanonical residues using placeholder symbols (“X”). To resolve these ambiguities, SMILES representations were retrieved based on therapeutic names when available and cross-validated against reported chemical formulas using PubChem [Kim et al., 2023].

Clear numerical entries were standardized by converting all measurements to seconds, for example, “0.43 hours” to 1548 seconds, and “3.8 ± 0.6 hrs” to 13,680 seconds using the central value. When multiple half-life measurements were reported for a single peptide under different experimental conditions, all entries were retained to reflect condition-dependent variability. The heterogeneity of half-life annotations reflects variability in experimental protocols rather than preprocessing artifacts. For PEPlife [Mathur et al., 2016], canonical SMILES and unambiguous compound identifiers were generally unavailable, and annotations frequently mixed D-amino acids and unnatural residues based on literature reports. Due to inconsistent or incomplete chemical specifications, explicit reconstruction of most unnatural amino acids was not attempted. Instead, peptides containing N–C cyclization or D-amino acids were treated as modified entries, while only non-cyclic, non-`X`-containing sequences with standard L-amino acid chirality were considered wild-type. D-amino-acid-containing SMILES were generated using the SMILES2PEPTIDE transformer from Tang et al. [2025c].

After cleaning, approximate SMILES construction, unit normalization, and duplicate removal based on exact matches of sequence and half-life duration, the final dataset contained 245 unique peptide entries, including both wild-type and chemically modified peptides. Among these, 130 entries correspond to amino acid–based peptide representations without explicit noncanonical modifications. For interpretability, all half-life values were converted from seconds to hours for reporting and evaluation.

## Supplementary Figures and Tables

**Figure S1:**
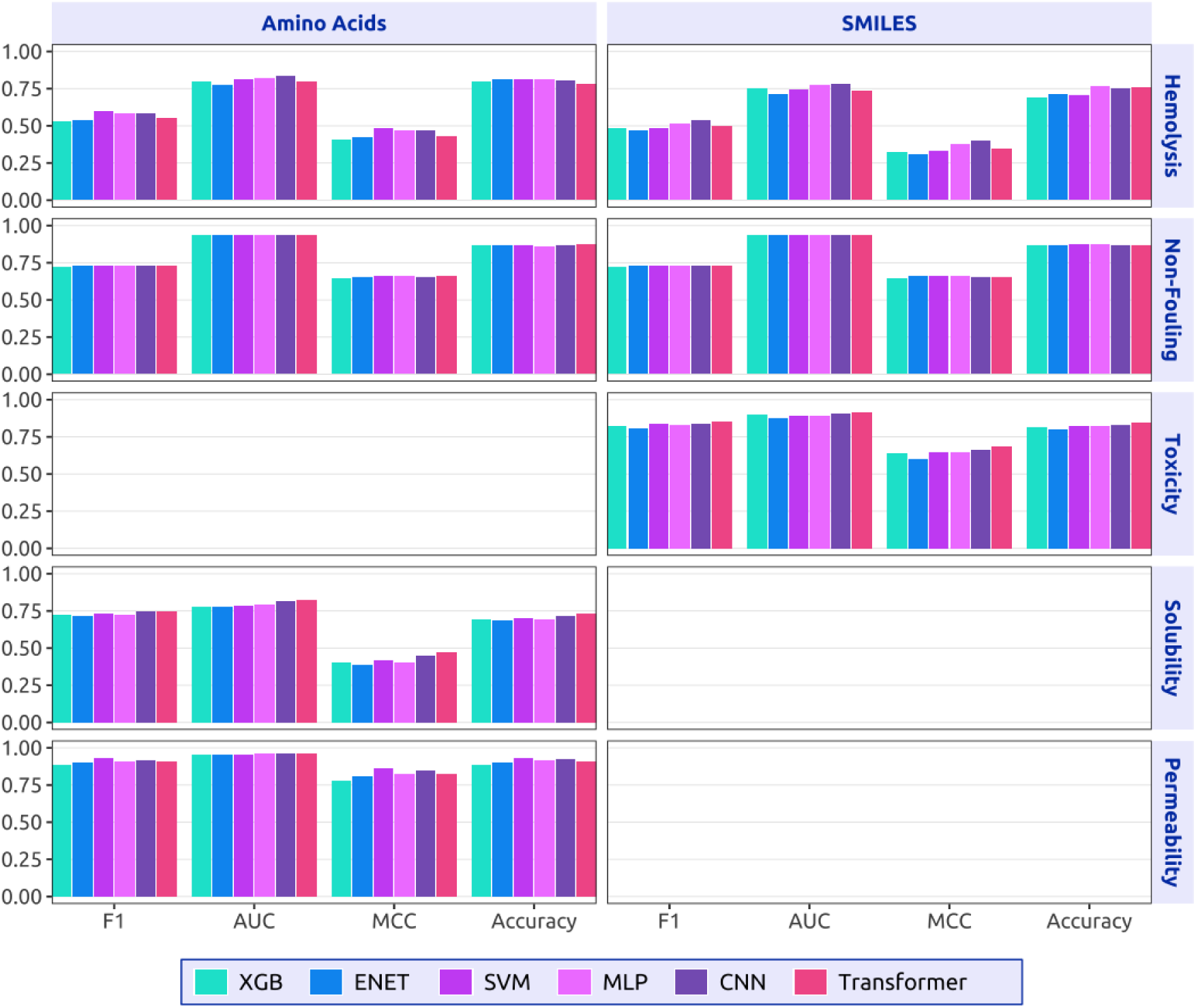
Per class performance across models. The best performing models were selected based on validation performance, refit on the validation set, and evaluated for final reporting. Properties without data are not reported.

**Figure S2:**
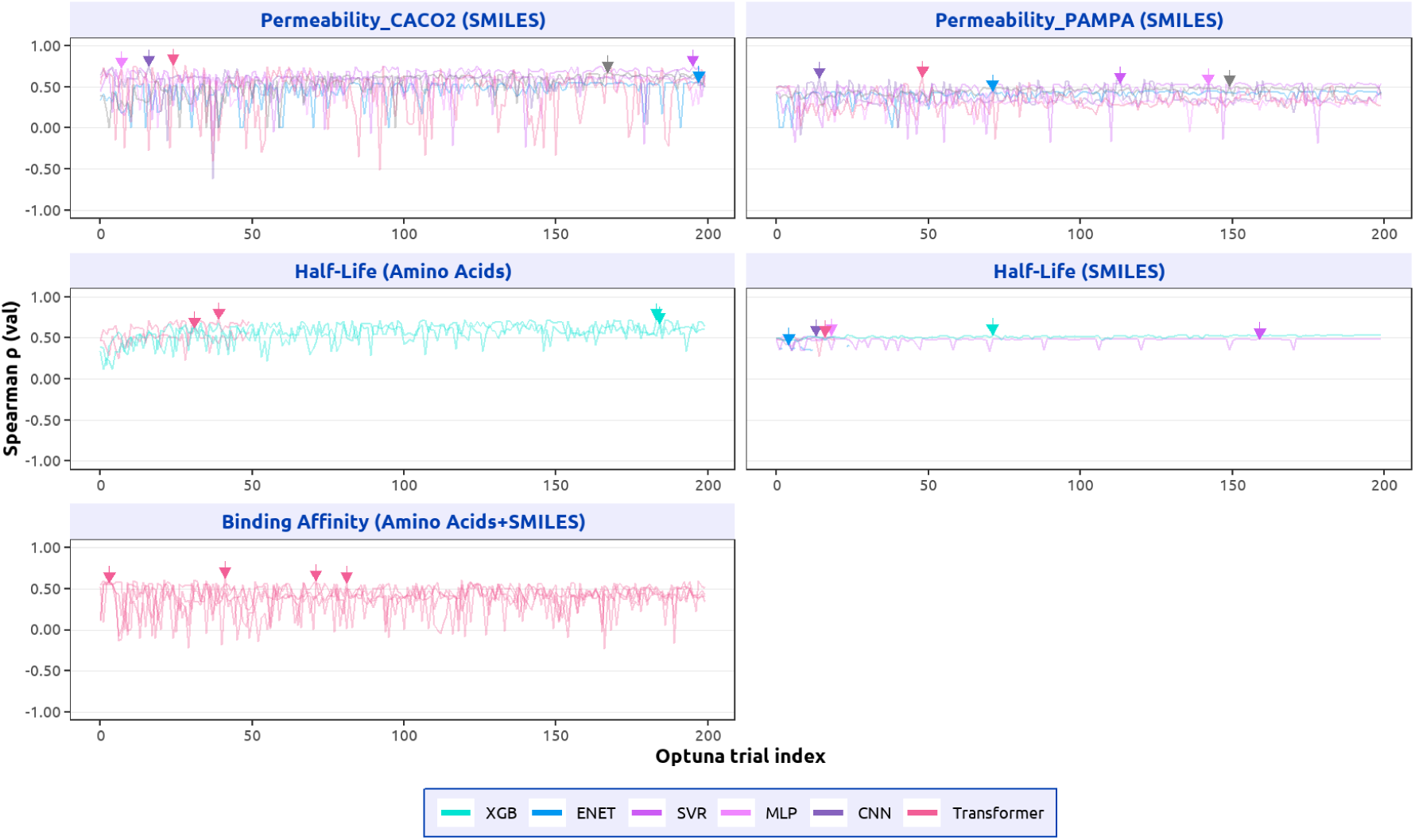
Optuna optimization traces for regression models. The arrow marks the selected trial with the best Spearman correlation (*ρ*) for each run. Machine-learning-based algorithms (XGB, ENET, SVR) were optimized for 200 trials, and neural-network-based algorithms (MLP, CNN, Transformers, where applicable) were optimized for 50 trials. SMILES tokenization results in longer input sequences and, therefore, slower training for neural-network-based models. The model choices for half-life and binding affinity reflect computational constraints and design considerations.

**Figure S3:**
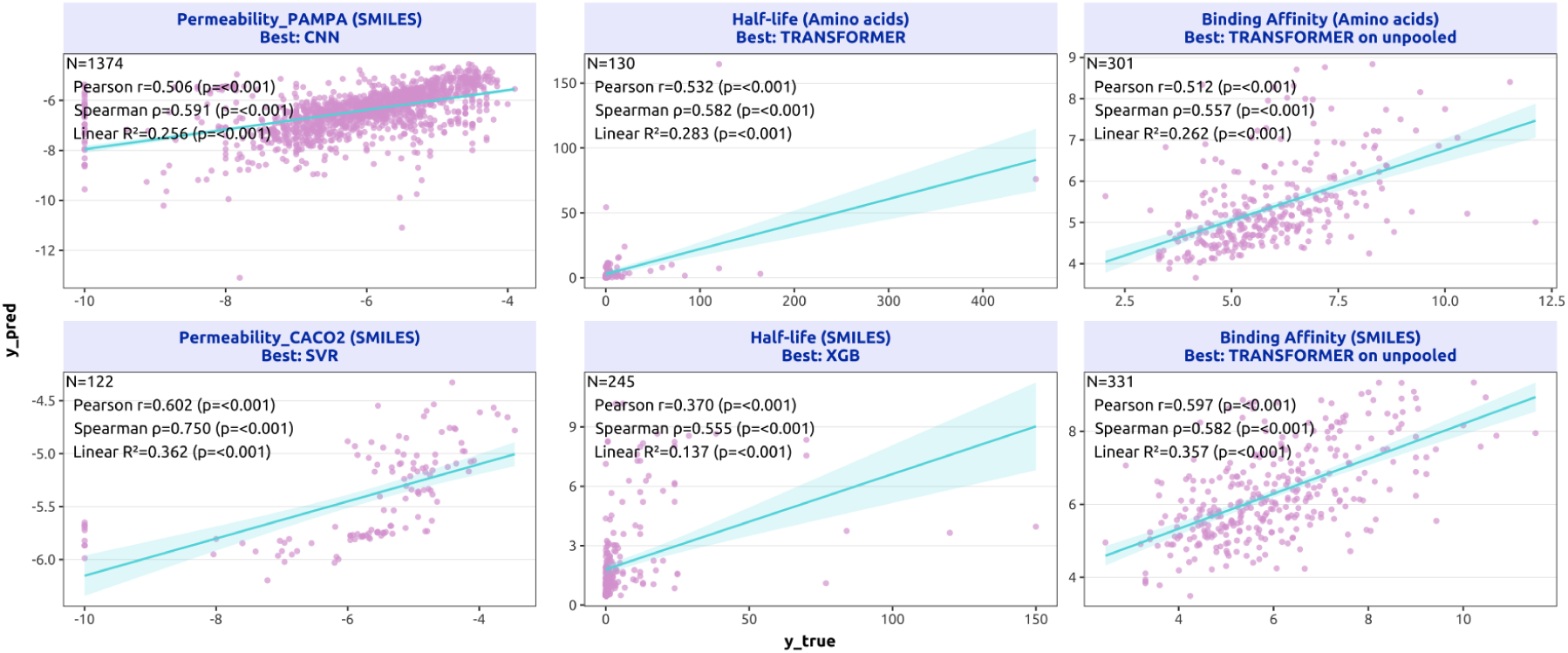
Top model performance for regression models. The best-performing models were selected and refit on the validation set for final evaluation. For half-life prediction, models were trained using cross-validation. The correlations are reported based on out-of-fold predictions collected during training. Specifically, in *k*-fold cross-validation, each fold’s predictions were generated by models trained on the remaining *k −* 1 folds, and the final results aggregate all held-out predictions.

**Figure S4:**
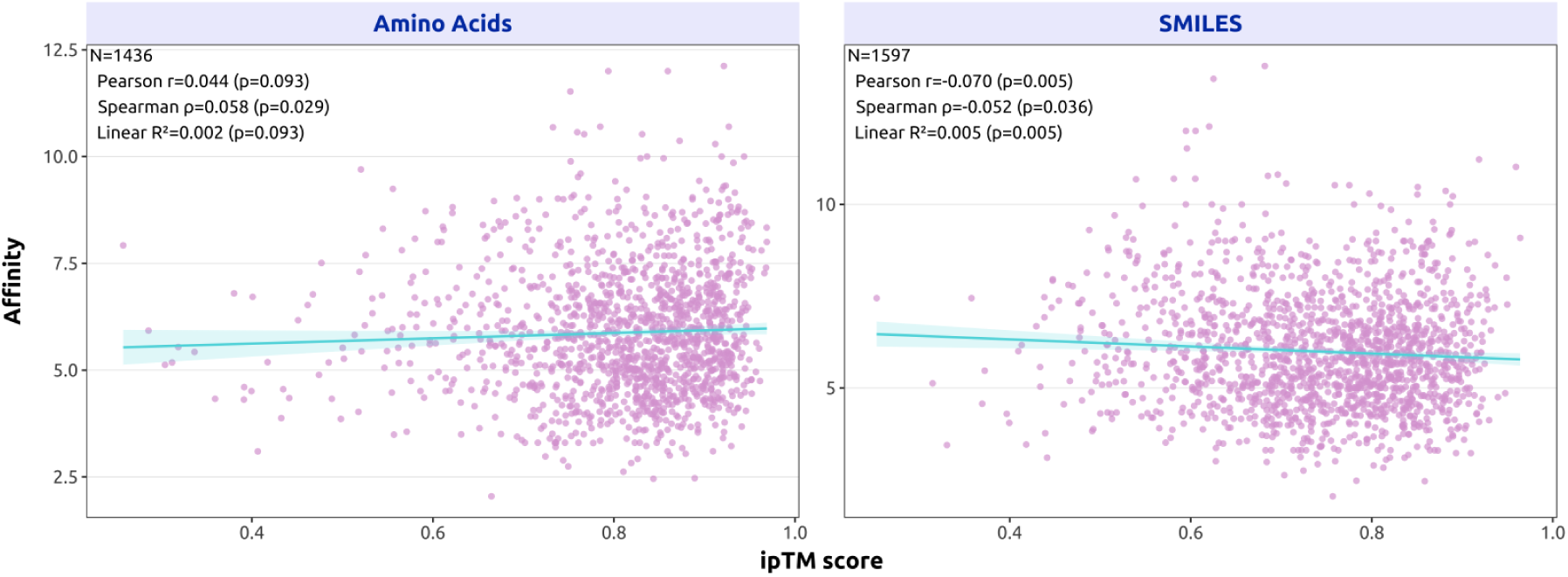
**Correlation between normalized experimental affinity score and Openfold3 ipTM score.**

**Figure S5:**
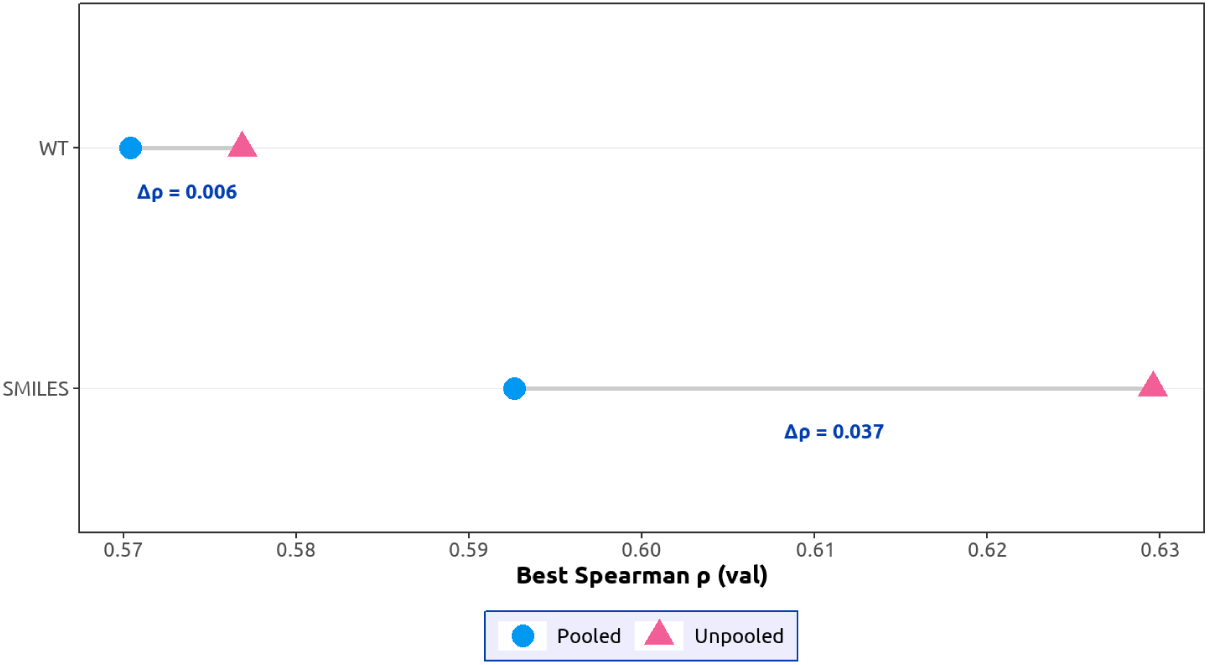
Unpooled versus pooled embeddings for binding affinity. Best validation Spearman *ρ* from Optuna optimization for WT and SMILES inputs are shown. The unpooled embeddings outperform pooled embeddings in both settings (*ρ* difference shown).

**Figure S6:**
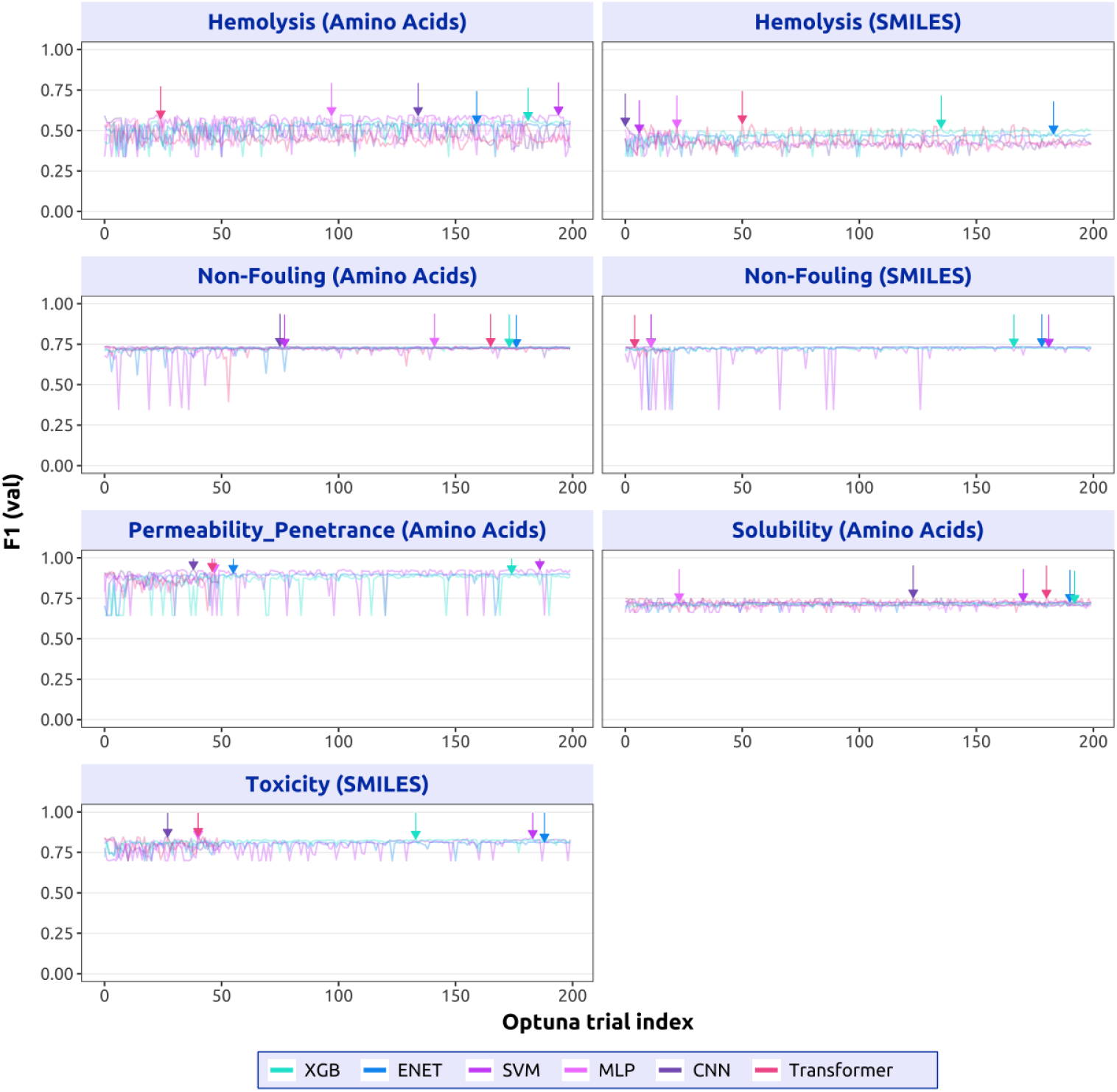
Optuna optimization traces. The arrow marks the selected trial with best F1 value for each run. All models were optimized for 200 trials.

**Table S1:** Final classification thresholds.

| Properties | Best Model |  | Classification Thresholds |  | Best F1 |  |
| --- | --- | --- | --- | --- | --- | --- |
|  | Amino acid | SMILES | Amino acid | SMILES | Amino acid | SMILES |
| Hemolysis | SVM | Transformer | 0.2521 | 0.4343 | 0.5975 | 0.5211 |
| Non-Fouling | MLP | ENET | 0.57 | 0.6969 | 0.7388 | 7356 |
| Solubility | CNN | - | 0.377 | - | 0.7537 | - |
| Permeability (Penetrance) | SVM | - | 0.5493 | - | 0.9291 | - |
| Toxicity | - | Transformer | - | 0.3401 | - | 0.8455 |

**Table S2:** XGBoost hyperparameters for classification and regression tasks. The ranges indicate Optuna search space boundaries. *^†^*Indicates log-uniform sampling in log space.

| XGBoost Hyperparameters |  |  |
| --- | --- | --- |
| Hyperparameter | Classification | Regression |
| Objective | binary:logistic | reg:squarederror |
| Eval metric | logloss | rmse |
| Lambda ($\lambda$) | $[10^{-8}, 50]^{\dagger}$ | $[10^{-10}, 10^2]^{\dagger}$ |
| Alpha ($\alpha$) | $[10^{-8}, 50]^{\dagger}$ | $[10^{-10}, 10^2]^{\dagger}$ |
| Gamma ($\gamma$) | $[0, 10]$ | $[0, 10]$ |
| Max depth | $[2, 15]$ | $[2, 16]$ |
| Min child weight | $[1, 500]$ | $[10^{-3}, 500]^{\dagger}$ |
| Subsample | $[0.5, 1.0]$ | $[0.5, 1.0]$ |
| Colsample by tree | $[0.3, 1.0]$ | $[0.3, 1.0]$ |
| Learning rate | $[10^{-3}, 0.3]^{\dagger}$ | $[10^{-3}, 0.3]^{\dagger}$ |
| Boosting rounds | $[50, 1500]$ | $[50, 2000]$ |
| Early stopping | $[20, 200]$ | $[20, 200]$ |
| Tree method | hist | hist |

**Table S3:** Elastic Net hyperparameters for classification and regression tasks. †Indicates log-uniform sampling. The class weight and selection are categorical choices.

| Elastic Net Hyperparameters |  |  |
| --- | --- | --- |
| Hyperparameter | Classification | Regression |
| Model type | cuLogReg, penalty="elasticnet" | cuElasticNet |
| Solver | qn (cuML) | cuML |
| $C$ (inverse regularization) | $[10^{-4}, 10^3]^{\dagger}$ | - |
| $\alpha$ (regularization) | - | $[10^{-8}, 10]^{\dagger}$ |
| $\ell_1$ ratio | $[0, 1]$ | $[0, 1]$ |
| Class weight | {balanced} | - |
| Selection | - | {cyclic, random} |
| Max iterations | $[200, 5000]$ | $[1000, 20000]$ |
| Tolerance | $[10^{-6}, 10^{-2}]^{\dagger}$ | $[10^{-6}, 10^{-2}]^{\dagger}$ |

**Table S4:**
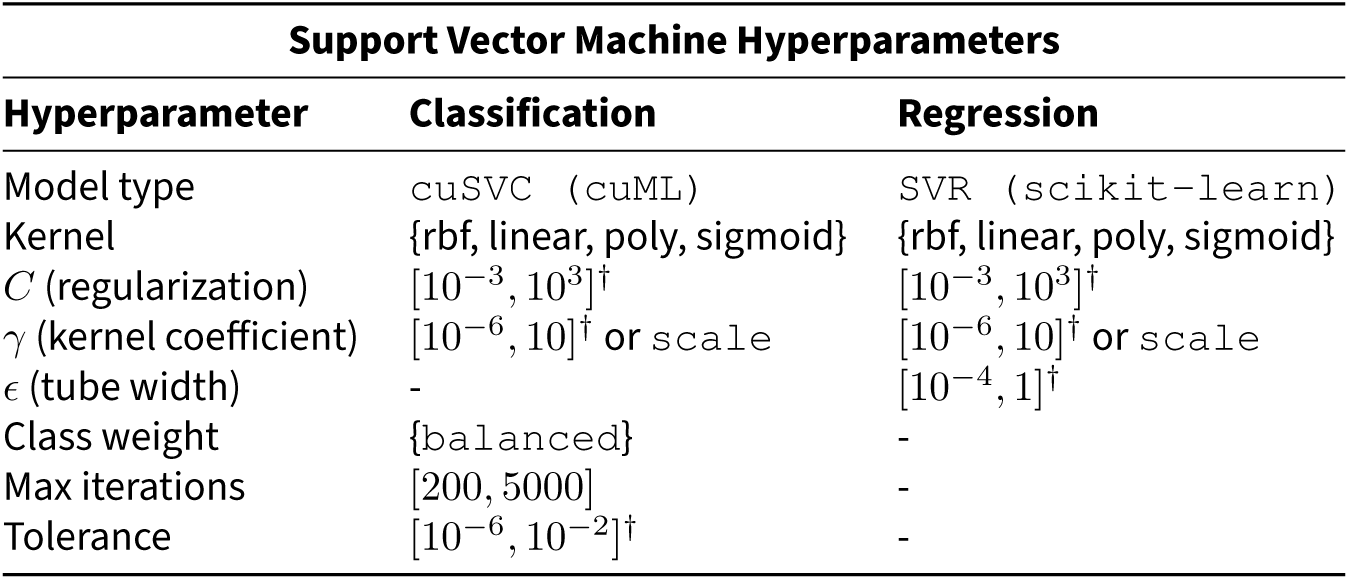
Support Vector Machine hyperparameters for classification and regression tasks. †Indicates log-uniform sampling. *γ* only applies to non-linear kernels.

**Table S5:**
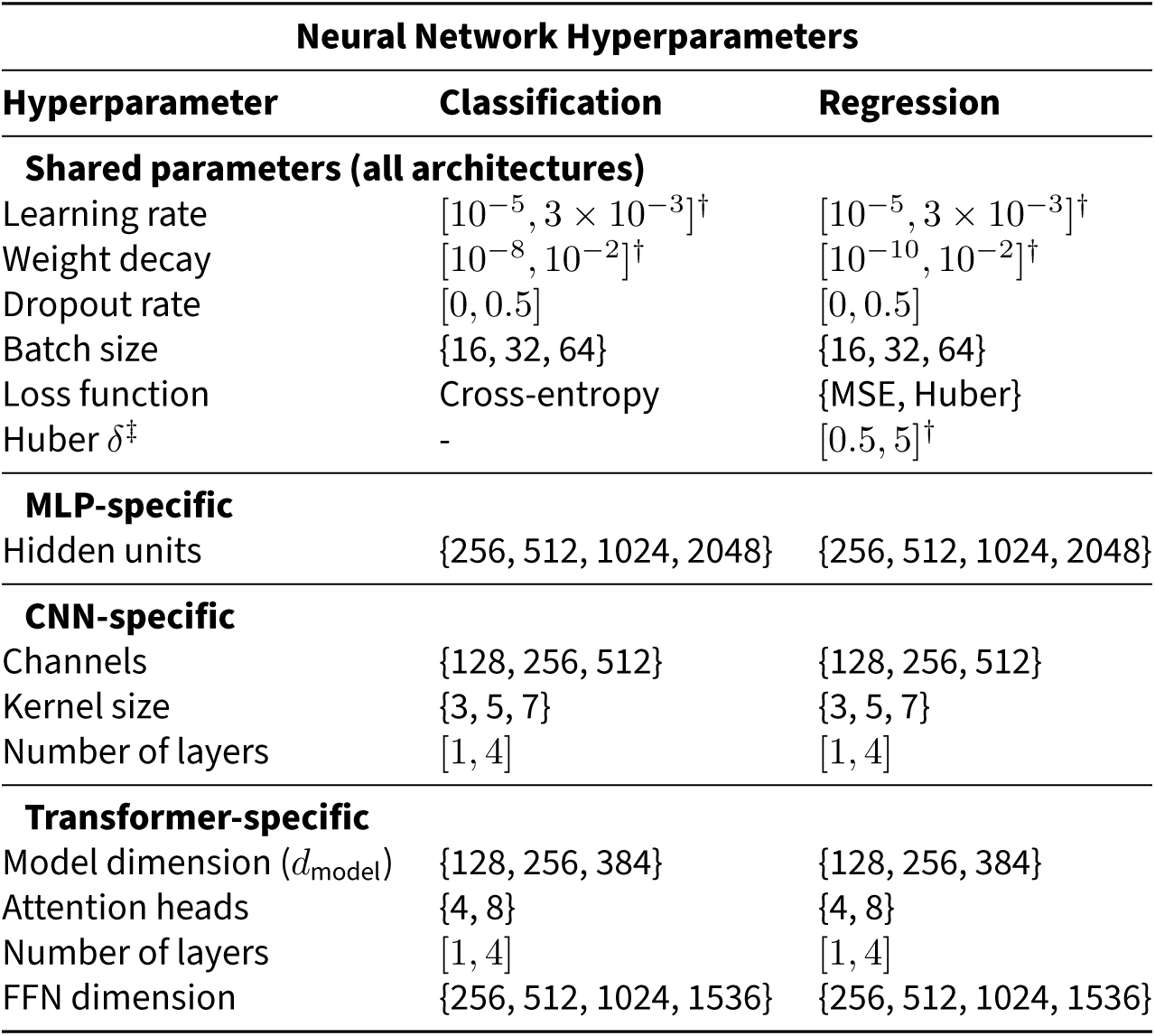
Neural network hyperparameters for classification and regression tasks. All models share the same base hyperparameters, with additional hyperparameters introduced only where required by the specific architecture. †Indicats log-uniform sampling. ‡The Huber δ controls outlier sensitivity.

| Neural Network Hyperparameters |  |  |
| --- | --- | --- |
| Hyperparameter | Classification | Regression |
| **Shared parameters (all architectures)** |  |  |
| Learning rate | $[10^{-5}, 3 \times 10^{-3}]^{\dagger}$ | $[10^{-5}, 3 \times 10^{-3}]^{\dagger}$ |
| Weight decay | $[10^{-8}, 10^{-2}]^{\dagger}$ | $[10^{-10}, 10^{-2}]^{\dagger}$ |
| Dropout rate | [0, 0.5] | [0, 0.5] |
| Batch size | {16, 32, 64} | {16, 32, 64} |
| Loss function | Cross-entropy | {MSE, Huber} |
| Huber $\delta^{\ddagger}$ | - | $[0.5, 5]^{\dagger}$ |
| **MLP-specific** |  |  |
| Hidden units | {256, 512, 1024, 2048} | {256, 512, 1024, 2048} |
| **CNN-specific** |  |  |
| Channels | {128, 256, 512} | {128, 256, 512} |
| Kernel size | {3, 5, 7} | {3, 5, 7} |
| Number of layers | [1, 4] | [1, 4] |
| **Transformer-specific** |  |  |
| Model dimension ($d_{\text{model}}$) | {128, 256, 384} | {128, 256, 384} |
| Attention heads | {4, 8} | {4, 8} |
| Number of layers | [1, 4] | [1, 4] |
| FFN dimension | {256, 512, 1024, 1536} | {256, 512, 1024, 1536} |

**Table S6:** Binding affinity model hyperparameters. A cross-attention architecture was used for peptide-protein binding prediction. †Indicats log-uniform sampling.

| **Binding Affinity Predictor Hyperparameters** |  |
| --- | --- |
| **Hyperparameter** | **Value / Range** |
| **Optimizer settings** |  |
| Learning rate (AdamW) | $[10^{-5}, 3 \times 10^{-3}]^{\dagger}$ |
| Weight decay | $[10^{-10}, 10^{-2}]^{\dagger}$ |
| **Architecture parameters** |  |
| Hidden dimension | {256, 384, 512, 768} |
| Attention heads | {4, 8} |
| Cross-attention layers | [1, 4] |
| Dropout rate | [0, 0.4] |
| **Training configuration** |  |
| Batch size | {16, 32, 64, 128} |
| Auxiliary loss weight ($\lambda_{\text{cls}}$) | $[0.1, 2.0]^{\dagger}$ |
| Primary loss | MSE (binding affinity) |
| Auxiliary loss | BCE (binding classification) |
| Max epochs | 50 |
| Early stopping patience | 10 |

